# Hsp40s display class-specific binding profiles, serving complementary roles in the prevention of tau amyloid formation

**DOI:** 10.1101/2021.04.11.439324

**Authors:** Rose Irwin, Ofrah Faust, Ivana Petrovic, Sharon Grayer Wolf, Hagen Hofmann, Rina Rosenzweig

## Abstract

The microtubule-associated protein, tau, is the major subunit of neurofibrillary tangles, forming insoluble, amyloid-type aggregates associated with neurodegenerative conditions, such as Alzheimer’s disease. Tau aggregation, however, can be prevented in the cell by a class of proteins known as molecular chaperones, which play important roles in maintaining protein homeostasis. While numerous chaperones are known to interact with tau, though, little is known about the detailed mechanisms by which these prevent tau aggregation. Here, we describe the effects of the ATP-independent Hsp40 chaperones, DNAJA2 and DNAJB1, on tau amyloid fiber formation and compare these to the well-studied small heat shock protein HSPB1. We find that each chaperone prevents tau aggregation differently, by interacting with distinct sets of tau species along the aggregation pathway and thereby affecting their incorporation into fibers. Whereas HSPB1 only binds tau monomers, DNAJB1 and DNAJA2 recognize aggregation-prone tau conformers and even mature fibers, thus efficiently preventing formation of tau amyloids. In addition, we find that both Hsp40s bind tau seeds and fibers via their C-terminal domain II (CTDII), with DNAJA2 being further capable of recognizing tau monomers by a second, different site in CTDI. These results provide important insight into the molecular mechanism by which the different members of the Hsp40 chaperone family counteract the formation, propagation, and toxicity of tau aggregates. Furthermore, our findings highlight the fact that chaperones from different families and different classes play distinct, but complementary roles in preventing pathological protein aggregation.

## INTRODUCTION

Tau is an intrinsically disordered protein (IDP) that is highly expressed in neurons and plays essential roles in microtubule self-assembly and stability (1), axonal transport (2), and neurite outgrowth (3). Tau binds to microtubules via its central microtubule-binding repeat (MTBR) domain (Fig. 1A), an interaction that is modulated by post-translational modifications (PTMs). Aberrant PTMs such as hyperphosphorylation and acetylation (4–6) were suggested to decrease the affinity of tau to microtubules, thus subsequently reducing microtubule stability. In addition, when tau dissociates from microtubules, it can form oligomers with the potential to disrupt cellular membranes, thereby impairing synaptic and mitochondrial functions, before ultimately forming amyloid fibers (7).

**Figure 1.**
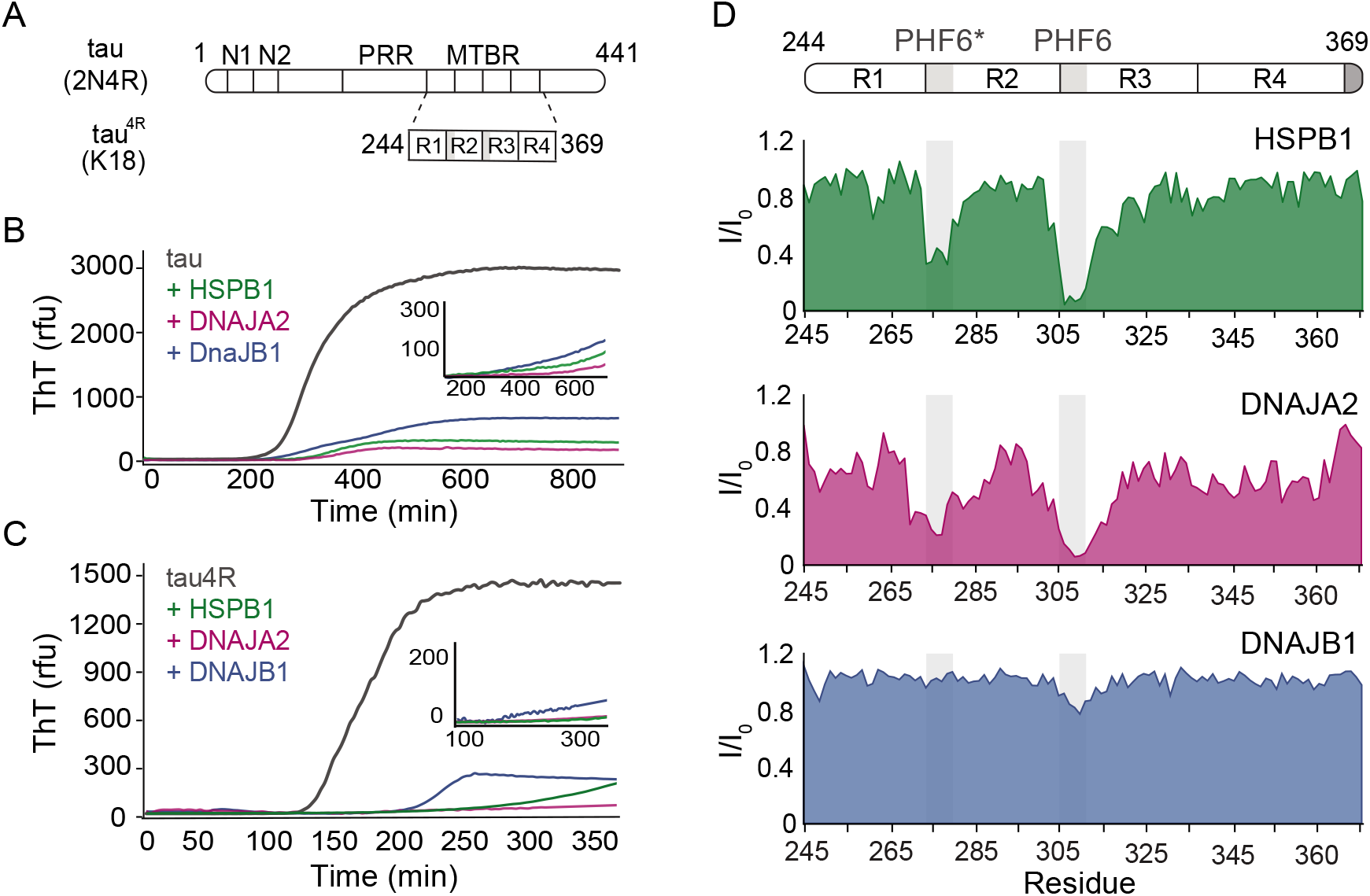
Interaction of chaperones with monomeric tau. **(A)** Domain organization of the longest splice isoform of tau protein (2N4R), and the short variant (4R / K18) containing only the microtubule binding repeats (MBTR). The location of the two N-terminal inserts (N1 and N2) and the polyproline region (PPR) is indicated. The MTBR region of Tau consists of four partially repeated sequences, R1 to R4, with the PHF6* and PHF6 aggregation-driving hexapeptides highlighted in grey. **(B-C)** ThT-based aggregation assay of 2N4R (B) and 4R (C) tau variants (10 μM) in the presence of 5 μM HSPB1 (green), DNAJA2 (purple), or DNAJB1 (blue) chaperones. The insert shows tau aggregation profiles in the presence of two-fold excess (20 μM) of the same chaperones, showing complete inhibition over the course of the experiment. Representative data from 3 independent experiments is shown. **(D)** Tau^4R^ binding profiles to HSPB1 (green), DNAJA2 (purple), and DNAJB1 (blue) chaperones, probed by NMR. Changes in NMR intensity ratios (I/I_0_) upon addition of two fold excess of each chaperone are plotted as a function of tau^4R^ residue number. The grey boxes represent the positions of the tau PHF6* and PHF6 aggregation-prone motifs. Values lower than 0.5 indicate intermolecular interactions.

These abnormal forms of tau are thought to play a key role in the pathogenesis of various human tauopathies, including Alzheimer’s disease (AD), frontotemporal dementias, and progressive supranuclear palsy (8). In such cases, tau forms large intracellular aggregates, termed neurofibrillary tangles, whose abundance and localization in the brain correlates with cognitive decline (8, 9). It is still unclear, however, if the fibrils themselves are the neurotoxic species, or whether prefibrillar soluble aggregates and oligomers of tau promote neuronal death by spreading tau pathogenicity from cell to cell in a prion-like manner.

The MTBR consists of four pseudo-repeats (R1-R4; Fig. 1A) of 30 residues each, with the 6-residue aggregation-prone regions located at the start of the second and third repeats. These two motifs, ^275^VQIINK^280^ and ^306^VQIVYK^311^, also called PHF6* and PHF6, respectively, enable the formation of β-sheet structures that are a prerequisite for tau aggregation (10, 11).

These aggregation-prone regions of tau are naturally also the primary binding site for members of the Hsp70 and Hsp90 chaperone families (12–14), which play an important role in tau homeostasis by regulating the interaction of tau with microtubules, counteracting its aggregation into amyloids, and targeting the misfolded species for degradation (15, 16). Mechanistically, the Hsp70 interaction with tau suppresses the formation of aggregation-prone tau nuclei as well as sequesters tau oligomers and fibrils (17, 18), thereby neutralizing their ability to damage membranes and seed further tau aggregation (19).

Other potent suppressors of tau aggregation are small heat shock proteins, and prominently amongst them, the HSPB1 chaperone. Like Hsp70 and Hsp90, HSPB1 interacts with the tau PHF6* and PHF6 regions (18, 20). However, its tau aggregation-prevention mechanism has been found to be markedly different. HSPB1 is a large oligomeric ATP-independent chaperone that was shown to delay tau fiber formation by weakly interacting with early species in the aggregation reaction (18). Interestingly, whereas HSPB1 impedes aggregation, it is unable to inhibit the elongation of fibrils once sufficiently large nuclei have formed (18).

Yet, HSPB1, is not the only ATP-independent chaperone reported to interfere with tau aggregation pathways. Hsp40s (also known as J-domain proteins, JDPs) are co-chaperones of the Hsp70 system that can also function as bona fide chaperones by preventing aggregation through their holdase activity and most likely also by modifying the conformation of client proteins (21). DNAJA2, a member of this JDP family, was recently identified as a potent suppressor of tau aggregation, capable of effectively preventing the seeding of tau and formation of amyloids in cells (13, 22). Additionally, another member of the JDP family, the class B DNAJB1, was recently shown to work with the Hsp70 system to disintegrate tau amyloid fibers extracted from AD brain tissues (23, 24).

Little is known, however, regarding how these co-chaperones interact with tau or the mechanism by which they modify tau disease-related amyloid states.

Intrigued by the possibility that the JDP chaperones may play an important role in tau homeostasis, we used NMR spectroscopy, in combination with kinetic aggregation assays, to elucidate the effect of the DNAJA2 and DNAJB1 chaperones on tau aggregation. Strikingly, we found that the aggregation-prevention mechanisms of DNAJB1 and DNAJA2 differ from that of HSPB1 holdase chaperone. Moreover, we found that the two Hsp40 family members also diverge in their interactions with tau - whereas DNAJA2 interacts with all species along the tau aggregation pathway, including inert tau monomers, DNAJB1 only interacts with aggregation-prone tau conformers, such as seeding competent species or mature fibers.

## RESULTS

### The chaperones slow down tau aggregation and bind tau

We first investigated the effect of DNAJA2 and DNAJB1 on the fibril formation of full-length tau and compared it to that of HspB1, a well characterized suppressor of tau aggregation. Tau aggregation was monitored using Thioflavin-T (ThT) fluorescence (25). The 3D GXG variant of HSPB1 (18) was used to mimic the fully activated dimeric form of the chaperone. As expected, the addition of HSPB1 significantly inhibited tau fiber formation, in agreement with previous reports (13, 18, 20). Interestingly, addition of DNAJA2 and DNAJB1 chaperones also completely inhibited the formation of tau fibrils for over 16 hours (Fig. 1B), with no observable ThT signal being detected over this length of time. Similar results were obtained with the shorter tau construct tau^4R^ (residues 244-372), which forms the core of tau filaments and nucleates tau aggregation (26). Despite the faster aggregation kinetics of tau^4R^ relative to the full-length tau (t½≈110 min vs t½≈340 min), the presence of even a sub-stoichiometric concentration of either of the 3 molecular chaperones (DNAJB1, HSPB1, or DNAJA2), fully suppressed aggregation for the duration of the experiment (10 hours) (Fig. 1C).

As all three chaperones are able to suppress tau aggregation, we next aimed to understand the mechanisms by which they do so.

In order to unravel the mechanism of aggregation prevention, we first identified the binding sites for the three chaperones on tau. To this end, we recorded ^1^H-^15^N HSQC spectra of either ^15^N-tau or ^15^N-tau^4R^ in the absence and presence of each chaperone. Upon addition of HSPB1, we observed significant peak broadening of tau^4R^ residues 275-280 and 306-311, which correspond to the PHF6* and PHF6 motifs (Fig. 1D and S1), in agreement with previous reports (13, 18, 20). Notably, the PHF6 motifs, which are the most hydrophobic regions within tau, are also the preferential binding sites for the major ATP-dependent chaperone families, such as Hsp70 and Hsp90 (13). A similar preference for the PHF6 and PHF6* motifs was also found for DNAJA2, with residues 275-284 and 306-320 showing significant peak broadening (Fig. 1D and S1B). Surprisingly, however, DNAJB1 showed no significant binding to tau^4R^, despite efficiently suppressing amyloid formation (Fig. 1D and S1B).

### Binding to tau fibrils

As DNAJB1 efficiently prevented tau aggregation in both full-length and tau^4R^ experiments, yet showed no detectable binding to the monomers, we hypothesized that it functions, instead, through association with preformed tau fibers. In fact, a similar behavior was recently reported for DNAJB1 in the case of α-synuclein, where the chaperone displayed a remarkable preference (>300-fold) towards the amyloid state of α-synuclein over the monomer (27).

We therefore checked whether DNAJB1 interacts with preformed tau^4R^ fibrils using co-sedimentation experiments. Indeed, a large portion of DNAJB1 was detected in the insoluble fraction together with tau^4R^ (Fig. 2A), indicating a strong interaction between tau fibrils and the chaperone. Co-sedimentation experiments with DNAJA2 and HSPB1 showed that also DNAJA2 co-precipitated with tau fibers whereas HSPB1 was mainly found in the soluble fraction and only marginally interacted with tau amyloids (Fig. 2A). Similar results were also obtained with fluorescence anisotropy, resulting in an affinity (dissociation constant) for tau^4R^ fibrils of 1.7 ± 0.2 μM (mean ± s.e.m.) for DNAJB1 and of 7.6 ± 0.6 μM for DNAJA2 (Fig 2B). In contrast, the affinity of HSPB1 for tau fibers was outside our experimental concentration range.

**Figure 2.**
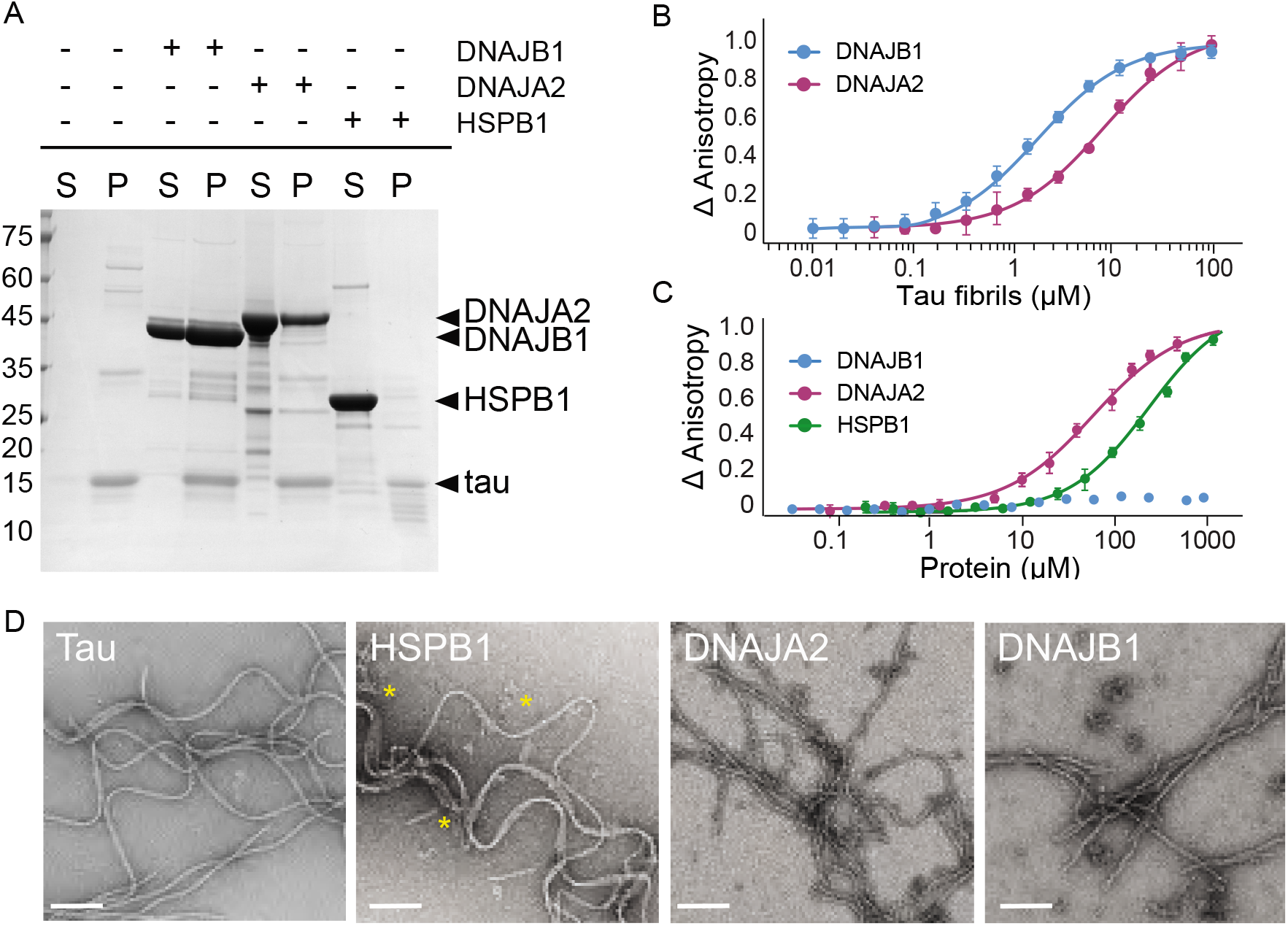
Interaction of chaperones with preformed tau fibrils. **(A)** Co-sedimentation of preformed tau amyloid fibrils (10 μM) in the presence of 10 μM DNAJB1, DNAJA2, or HSPB1 chaperones. SDS-PAGE of the supernatant (S) and pellet (P) fractions following ultracentrifugation is shown. **(B)** Fluorescence anisotropy assays of tau fibers binding to DNAJB1 (blue) or DNAJA2 (purple). The K_D_s were calculated as 1.7 ±0.2 μM and 7.6 ± 0.6 μM, respectively. **(C)** Fluorescence anisotropy assays of monomeric tau binding to HSPB1 (green), DNAJB1 (blue), or DNAJA2 (purple). The K_D_s were calculated as 250 ± 26 μM for HSPB1 and 43 ± 8 μM for DNAJA2. No binding was observed for DNAJB1. **(D)** Representative negative stain EM micrographs of preformed tau fibers alone or upon incubation with 1:1 molar ratio of HSPB1, DNAJA2, or DNAJB1 chaperones. The asterisks show the position of HSPB1 chaperone clusters. White bar is 200 nm.

The selective interaction of DNAJB1 and DNAJA2 chaperones with tau fibers was also observed by negative stain electron microscopy (EM). Here, tau alone formed characteristic paired helical filaments with a periodicity of 50-100 nm (Fig. 2D), consistent with previous observations (28). Upon addition of HSPB1 chaperone to the fibers, no changes to fiber length or morphology were detected, in agreement with our co-sedimentation assays that showed no binding of HSPB1 chaperone to the preformed fibers. HSPB1 itself, however, generated large protein assemblies that can be seen next to the fibers in the EM images (marked by *).

Addition of DNAJB1 or DNAJA2, on the other hand, significantly changed the fiber morphology to straight filaments, decorated by periodically-bound chaperones (Fig. 2D). The overall length of the fibers, however, did not change substantially, and also smaller tau fragments were not observed in our EM images (see fig. S2 for more representative images), indicating that DNAJB1 and DNAJA2 do not enhance fiber breakage or fragmentation upon binding. In summary, HSPB1 only binds tau^4R^ monomers, DNAJB1 interacts with tau^4R^ fibrils, whereas DNAJA2 binds both monomers and fibers.

### Each chaperone suppresses a specific subset of microscopic processes in the tau aggregation reaction

We were next interested to see how the different tau binding modes of the three chaperones affect their respective aggregation prevention mechanisms. We therefore performed a series of aggregation kinetic experiments, varying the concentration of the chaperones while maintaining a constant concentration of tau^4R^. In the absence of chaperones, the aggregation of tau^4R^ has previously been reported to occur through the following microscopic steps: primary nucleation, fibril growth through the addition and rearrangement of monomers (saturating elongation), and fiber fragmentation (29, 30) (Fig. S3). Indeed, an analysis of the half-saturation times (half-times) as a function of the tau^4R^ concentration shows a slight positive curvature, which is indicative of such a mechanism (31) (Fig. S3A). On average, the scaling exponent was determined to be −0.34 ± 0.04, which is consistent with a dominant primary nucleation pathway and a contribution stemming from the presence of fibril fragmentation (29, 30, 32) (see methods for more detail). Assuming a nucleus size of two tau^4R^ monomers (29, 30), a global fit of the kinetics of tau^4R^ in the absence and presence of aggregation seeds provided the kinetic rates for nucleation, elongation, and fragmentation, as well as the saturation constant (Fig. S3).

All chaperones caused a concentration-dependent retardation of tau aggregation at sub-stoichiometric concentrations (Fig. 3). To understand the effect of the chaperones at the microscopic level, we fit the data to the kinetic model of tau aggregation using the kinetic parameters determined in the absence of chaperone. In order to determine which step in the aggregation mechanism is most likely affected by each of the chaperones, we allowed individual kinetic rates to vary in the analyses (see methods for detail). Furthermore, as the addition of any of the chaperones to preformed tau fibrils did not change the overall fibril length (Fig. 2B and S2), we assume that the chaperones have no effect on tau fragmentation rates (k_m_).

**Figure 3.**
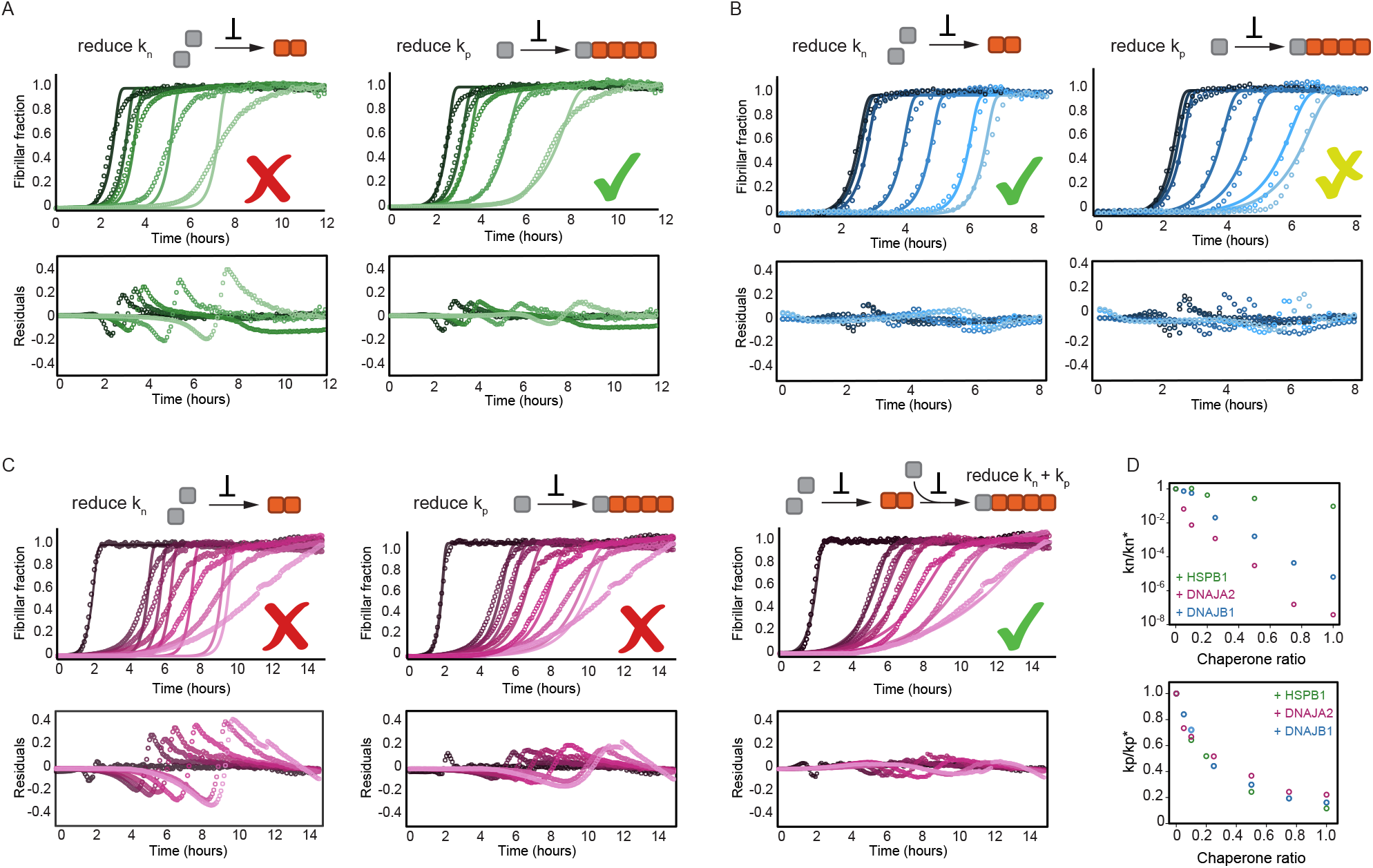
HSPB1, DNAJA2, and DNAJB1 chaperones affect different microscopic steps in the aggregation process. Kinetic profiles of the aggregation of 10 μM of tau in the absence (black) and presence of increasing concentrations of **(A)** HSPB1 chaperone (1, 2.5, 5, and 10 μM; dark to light green), **(B)** DNAJB1 (0.5, 1, 2.5, 5, 7.5 and 10 μM; dark to light blue), or **(C)** DNAJA2 (0.5, 1, 2.5, 5, 7.5 and 10 μM; dark to light purple). Open circles represent experimental data, and solid lines represent the fit of the kinetic profiles where only the primary nucleation (k_n_, left) or elongation (k_p_, right) pathways are inhibited. Residuals of the fits are shown under each panel. The changes in the aggregation kinetics caused by HSPB1 fit well with the elongation rate being primarily affected by the chaperone (A, right) whereas the changes in tau aggregation kinetics caused by DNAJB1 can be best described by the reduction of primary nucleation rates (B, left). In the case of DNAJA2, the changes in aggregation kinetics cannot be well described by the delay of only primary nucleation (left) or only elongation (middle) rates. There is good agreement, however, between the experimental data and fits to the integrated rate law, in which both primary nucleation and elongation events have been considered simultaneously. **(D)** The changes in microscopic nucleation (top) and elongation (bottom) rate constants as a function of the concentration of the molecular chaperones, relative to tau alone.

Upon fitting HSPB1 aggregation prevention data, only a poor agreement was achieved when allowing the perturbation of primary nucleation rates (k_n_) (Fig. 3A). Given the fact that HSPB1 binds tau monomers, the relatively moderate effect on tau primary nucleation was somewhat surprising as monomer binding should inhibit both nucleation and elongation. In contrast, fitting the fiber elongation rate (k_p_) provided a significantly better description of the kinetic data (Fig. 3A), indicating an order of magnitude reduction of this rate (Fig. 3D). Hence, we conclude that HSPB1 inhibits tau aggregation primarily by preventing the incorporation of tau monomers into the elongating fibers.

The effect of DNAJA2 on tau aggregation could neither be described by the reduction of nucleation rates nor of elongation rates alone (Fig. 3B). These results were not entirely surprising given that the chaperone can bind to both monomeric tau and fibrils (Fig. 1D and 2), This, then provides DNAJA2 at least two distinct pathways to impact aggregation, namely by (i) lowering the amount of monomeric tau accessible for nucleation and elongation and (ii) lowering the potency of fibrils to grow. Indeed, allowing the variation of both elongation and nucleation parameters achieved the best agreement with our data (Fig 3C), revealing substantial perturbation of both rates by DnaJA2 (Fig. 3D) and indicating that this chaperone affects the aggregation process in more intricate ways than HSPB1.

Surprisingly, the effect of DNAJB1 on tau aggregation was best described by its ability to reduce the rates of primary nucleation (Fig. 3B, left) and only to a lesser extent, fiber elongation (Fig. 3B, right). This ability of the DNAJB1 chaperone to effectively inhibit the rate of tau^4R^ primary nucleation (Fig. 3B) was unexpected as primary nucleation involves interaction between tau monomers. Yet, no interaction between tau^4R^ monomers and DNAJB1 was observed in our NMR experiments.

### Two conformations of monomeric tau

The effect of DNAJB1 on tau^4R^ nucleation, despite its inability to interact with monomeric tau^4R^, can, however, be reconciled by the recent finding that soluble monomeric tau^4R^ exist in two conformational ensembles – an ensemble that does not spontaneously aggregate (“inert” tau monomer) and a seed-competent monomer that triggers the spontaneous aggregation of tau (33, 34). It is therefore possible that DNAJB1 only identifies and interacts with the “aggregation-prone” tau species and not the inert monomers.

Such seed-competent species have been proposed to be readily populated in tauopathy-associated tau mutants (34), and can be generated *in vitro* by addition of polyanions such as heparin (35). Unfortunately, the addition of heparin causes the rapid formation of tau fibers, thus preventing a detailed structural investigation of the seed-competent ensemble using NMR-spectroscopy.

Yet, whereas tau^4R^ is inaccessible under conditions at which it rapidly aggregates, the rate of tau fibrillization can be tuned by altering the concentration of heparin present in the aggregation reaction. Low concentrations of heparin enhance the rate of fiber formation, whereas higher heparin concentrations potentially inhibit it (36). We therefore performed aggregation kinetics at different heparin concentrations (0.1 - 40 μM) to identify the conditions at which tau aggregation is sufficiently slow to permit NMR experiments. We found that heparin increased the rate of fiber formation up to a sub-stoichiometric concentration of 1 μM (1:0.1 tau:heparin) (Fig. S4A). Above this threshold, further addition of heparin in fact slowed tau aggregation in a dose dependent manner. At a 1-4 fold excess of heparin (10-40 μM), we found that tau amyloid formation was arrested for over two hours (Fig. S4A). An equimolar concentration of heparin to tau was therefore used to generate a soluble, aggregation-prone tau species (33) that does not aggregate and is therefore amenable for NMR experiments.

We then monitored chaperone binding to this tau species by recording ^1^H-^15^N HSQC spectra for ^15^N-tau-heparin complex alone, and upon addition of DNAJB1, DNAJA2, and HSPB1 chaperones to the mixture (Fig. 4). To map the binding sites for the chaperones on this heparin-bound, aggregation-prone tau species, we first had to assign its spectrum. HNCA, CBCA(CO)NH, HN(CA)CO, and HNCO 3D NMR experiments were recorded and assignments were obtained for 88% of the non-proline residues. Heparin binding resulted in chemical shift perturbations (CSPs) to the NMR spectrum of tau^4R^, mainly in the R1, R2 and PHF6* regions (Fig. 4A and S4B). No changes were observed to the overall dispersion of the spectrum, indicating that heparin binding does not induce global folding of tau, in agreement with previous findings (11).

**Figure 4.**
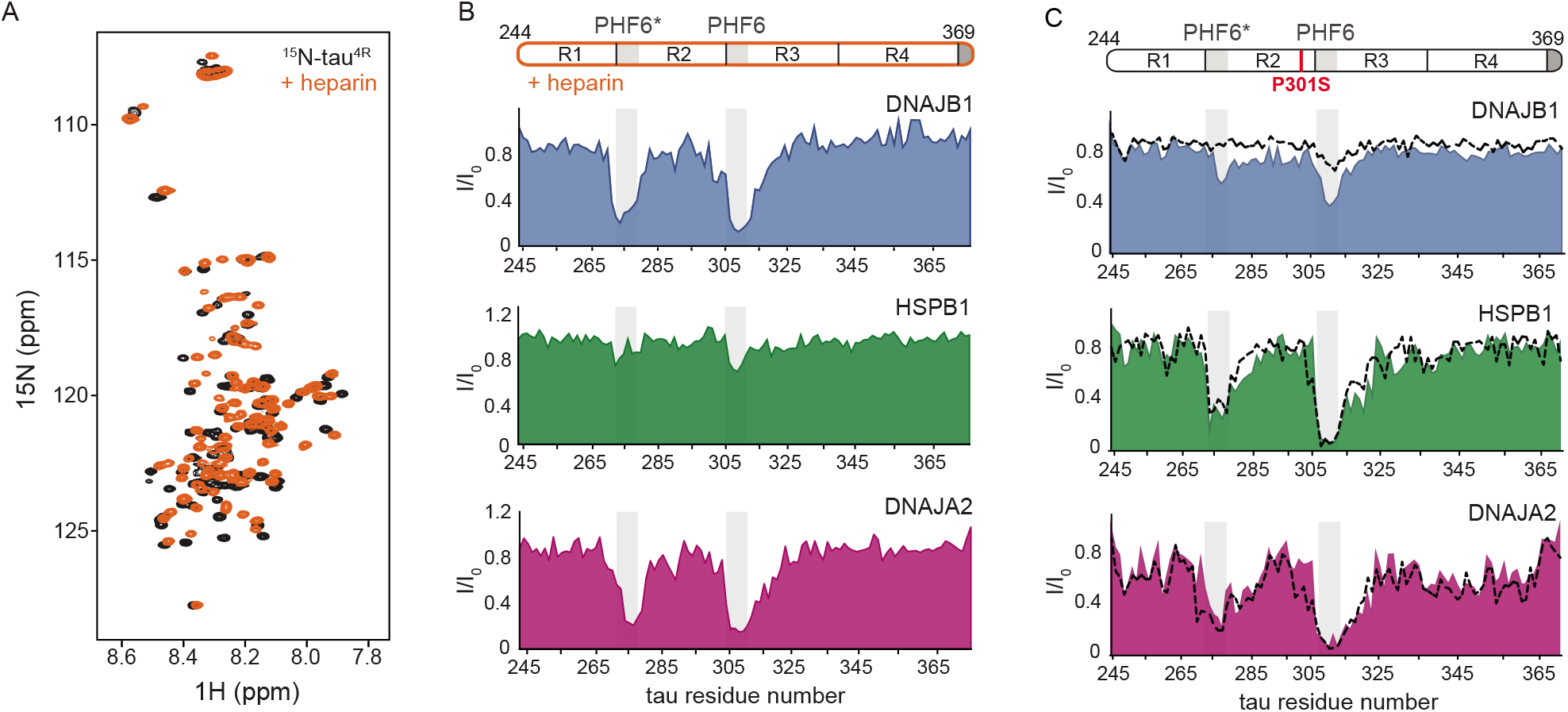
Interaction of chaperones with aggregation-prone tau species. **(A)** Overlay of two-dimensional ^1^H-^15^N NMR spectra of tau in the absence (black) and presence of heparin (orange). Heparin binds mainly to the R1, R2 and PHF6* regions (see S4C), with no changes observed to the overall dispersion of the spectra, indicating that heparin binding does not result in global folding of tau. **(B)** Residue-resolved NMR signal attenuation (I/I_0_) of tau in complex with heparin upon addition of one molar equivalent of DNAJB1 (blue), DNAJA2 (purple), or two molar equivalents of HSPB1 (green). Grey boxes represent the positions of tau PHF6* and PHF6 aggregation-prone motifs in tau. Unlike in the case of tau monomer, DNAJB1 interacts strongly with this aggregation-prone species, while HSPB1 chaperone does not. **(C)** Residue-resolved backbone amide NMR signal attenuation (I/I_0_) of tau P301S mutant upon addition of two molar equivalents of DNAJB1 (blue), HSPB1 (green), or DNAJA2 (purple) chaperones. Dashed lines represent the same intensity plots for wild-type tau (as in Figure 1D). Compared to the wild-type, P301S mutant shows increased interaction with DNAJB1.

With these assignments in hand we were able to identify that, despite having negligible affinity towards the tau monomer, the DNAJB1 chaperone indeed interacts strongly with the aggregation-prone tau-heparin mixture (Fig. 4B), as we previously predicted. We further mapped this binding to tau residues 275-280 and 305-314 of the PHF6 and PHF6* repeats - the same regions to which both DNAJA2 and HSPB1 bind in the inert, monomeric tau.

We next tested whether DNAJA2 and HSPB1 interact with the heparin-bound tau. DNAJA2 showed strong binding to the R2 and R3 PHF6* repeats of this aggregation-prone form of tau^4R^, similarly to its interaction with the free monomer (Fig. 4B). In contrast, HSPB1 only bound this species weakly, despite previously displaying a strong interaction with free tau^4R^. The lack of interaction between HSPB1 and aggregation-prone tau species explains our previous observation that HSPB1 does not affect the rate of fiber nucleation (Fig. 3A), as such an inhibition would require a direct interaction with the aggregate nucleus or aggregation-prone tau.

Thus, while interacting with the same regions of tau, the 3 chaperones each display specific preferences for the different tau monomer conformers: HSPB1 interacts solely with tau monomers; DNAJB1 exclusively with the aggregation-prone species, and DNAJA2 with both.

It was unclear, however, how the chaperones discriminate between the inert form of tau and the aggregation-prone tau-heparin complex.

### Aggregation prone tau species

Secondary-structure propensity analysis of free and heparin-bound tau demonstrated that the different chaperone binding profiles cannot be explained by heparin-induced formation of extended β-strands structures in the PHF6 repeats (35), as these regions displayed no increase in secondary-structure propensity upon addition of heparin (Fig. S4C).

We then set out to determine whether the differences between monomeric tau^4R^ and the heparin-tau complex are caused by electrostatic interactions. Heparin is a polyanion with a net charge of −3 per disaccharide at pH 7.0 (or estimated charge density of −6.7 e-/kD (37). In contrast, tau^4R^ has a positive net charge of +9.5, suggesting that the heparin-tau^4R^ complex is significantly stabilized by electrostatic attractions. As such, the ability of the chaperones to distinguish between the monomeric and aggregation-prone tau species may be related to simple difference in the net charge of the complex, with heparin binding, for example, reducing electrostatic repulsions that may exist between DNAJB1 and monomeric tau^4R^. Screening any such electrostatic interactions with higher ionic strengths (0.3 M), however, neither facilitated binding between DNAJB1 and tau^4R^ (Fig. S5) nor altered tau^4R^ interaction with HSPB1 or DNAJA2 chaperones (compare Fig. 1D and S5). Similarly, the interaction cannot be explained by attractive electrostatic interactions caused by the negative net-charge of the complex, as we did not detect specific binding between heparin and any of the chaperones. Hence, electrostatic interactions cannot explain the different binding profiles of the chaperones to the two tau species.

A clue to how heparin alters the conformational ensemble of tau^4R^ came from a recent structural characterization of patient-derived, seeding-competent tau monomer. This species of tau was reported to have an expanded ensemble with a more exposed PHF6 motif compared to the inert monomeric protein (33, 34). This increased exposure of the PHF6 aggregation motifs was, in turn, suggested to drive the self-assembly and subsequent aggregation of tau (38). Interestingly, binding of heparin to tau was also shown to expand the local conformation of the repeat regions (R2 and R3), thereby making the amyloidogenic PHF6 sequences more accessible (35). The increased accessibility of the PHF6 motif in the heparin-bound state may thus enable the interaction of DNAJB1 with tau.

To test whether the expansion of the seeding-competent tau species and the exposure of the PHF6 repeat are indeed the discriminating factors for chaperone binding, we monitored the interaction of the three chaperones with P301L/S missense mutations, which are known to cause dominantly inherited tauopathy (39). These tau variants had been shown to display more exposed PHF6 sequences (34) and could therefore mimic the aggregation-prone tau without requiring the addition of heparin. Similarly to what we observed for wild-type tau^4R^, we found that indeed all 3 chaperones efficiently suppressed the aggregation of these tau mutants (Fig. S6B), although with reduced efficacy in the case of HSPB1 and higher activity of DNAJB1 (Fig. S6C).

The interaction of the two familial mutations of tau, P301L and P301S, with HSPB1, DNAJB1, or DNAJA2 chaperones, were then assayed using NMR spectroscopy. The resulting binding profiles of the two tau variants in the presence of HSPB1 and DNAJA2 were very similar to those of wild-type tau, with residues 274-280 and 305-318, corresponding to PHF6 and PHF6* repeats, displaying severe peak broadening (Fig. 4C and S6). DNAJB1, on the other hand, while showing no binding to wild-type tau, caused substantial reductions in peak intensities in both PHF6 motifs of the P301S and P301L tauopathy mutants (Fig. 4C and S6A).

Thus DNAJB1-binding to monomeric tau indeed appears to depend on the exposure of the PHF6 region, thus allowing it to distinguish between inert tau and the aggregation-prone species that eventually leads to amyloid formation.

### Changes to tau fibers caused by the chaperones

Our results show that all three chaperones efficiently inhibit tau aggregation via interactions with the hydrophobic aggregation-prone PHF6 repeats. Each of the chaperones interacts with a specific set of tau species, thus slowing down different microscopic processes in the aggregation reaction. However, it remained unclear whether the chaperones also affect the size and/or morphology of tau fibers.

Since ThT fluorescence only reports on total fibril mass, with no differentiation to fibril length or number, we turned to EM in order to image the fibers. When adding HSPB1 at the start of the aggregation reaction, a clear dose-dependent decrease in the density of the tau fibrils was observed (Fig. 5A). Only a few fibers per micrograph were seen when in the presence of a 2-fold excess of HSPB1 chaperone which, interestingly, were significantly straighter in appearance compared to the paired-helical fibers of tau alone. The length of the formed fibers, however, remained unaltered (3-4 μm), regardless of the concentration of HSPB1 added. HSPB1 thus appears to only affect the number of fibrils, not the fibril length itself, indicating that its interaction with tau monomers merely slows the fibril elongation rates, but does not prevent their incorporation into the amyloid fibers. Moreover, this observation likewise suggests that HSPB1 binding to fiber ends, if present, is not sufficient to stop fibril growth once started.

**Figure 5.**
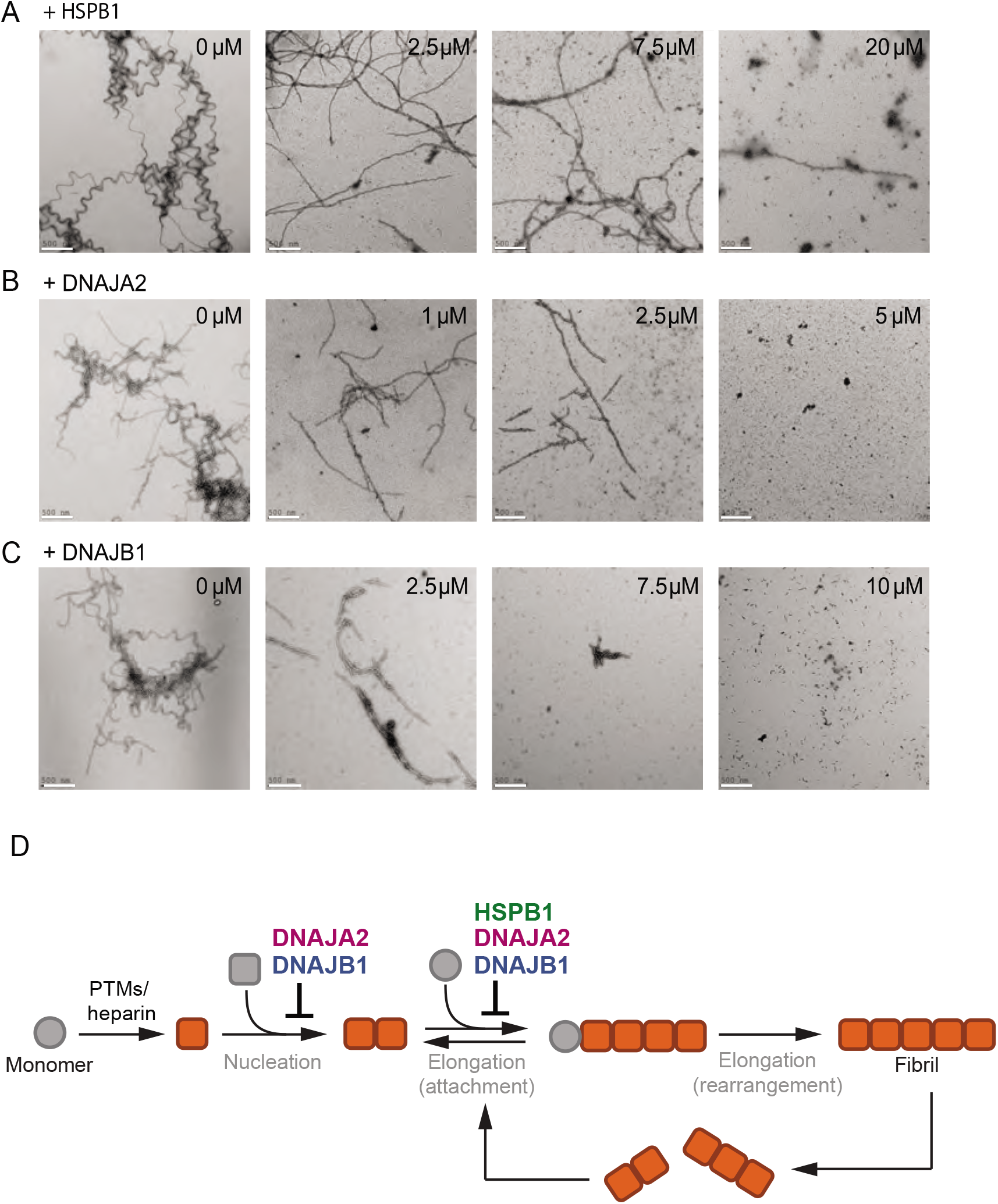
The chaperones change the morphology of tau amyloids. **(A-C)** Representative negative stain electron micrographs of end-products of the tau aggregationprevention assays preformed in the presence of indicated concentrations of HSPB1 (A), DNAJA2 (B) and DNAJB1 (C). White bar is 500 nm. Tau^4R^ forms long fibrils in the absence of chaperones and is still able to form long, fully developed fibrils in the presence of HSPB1 dimer. It forms smaller oligomeric species in the presence of both DNAJA2 and DNAJB1, indicating that these chaperones maintain much of the protein in a non-fibrillary state. **(D)** Overview of the variety of diverse microscopic mechanisms through which HSPB1, DNAJB1, and DNAJA2 molecular chaperones can suppress tau amyloid formation, as revealed by the binding and kinetic assays in this paper. These results demonstrate that the chaperones have evolved to exploit the different opportunities to modulate tau aggregation, by binding to different tau species along the aggregation pathway.

DNAJA2 and DNAJB1 chaperones, on the other hand, affected fiber formation differently when added to the tau aggregation reaction. In contrast to HSPB1, DNAJA2- or DNAJB1-containing reactions yielded shorter fibrils that further shortened with increasing chaperone concentrations (Fig. 5B and C). This result is not unexpected given that these two chaperones also bind to tau^4R^ fibers (Fig. 2), which likely contributed to the arrest of tau amyloid elongation.

Interestingly, DNAJA2 was very efficient in reducing both the size and amount of formed tau fibers, and even at the sub-stoichiometric concentration of 1:0.1 tau:DNAJA2, a significant reduction in fiber length was observed. At a ratio of 1:0.25 tau:DNAJA2, the majority of fibers were between 0.5-2.0 μm long, and at 1:0.5 tau:DNAJA2 ratio, no fibers were detected (Fig. 5B). Thus, the ability of DNAJA2 chaperone to interact with all tau species and to prevent both elongation and nucleation rates, makes it a potent suppressor of tau aggregation. DNAJB1, that does not interact with tau monomers, was less effective than DNAJA2 in reducing fiber size, and at 1:0.5 tau:DNAJB1 the fibers were still visible. These were, however, significantly shortened, with sizes ranging from 0.2-1.0 μm, and at a stoichiometric ratio of 1:1 no fibers were observed (Fig. 5C).

Thus, DNAJB1 and DNAJA2 chaperones, which bind to both aggregation-prone tau species and fibers, are significantly more efficient in preventing the elongation of tau into mature fibers than HSPB1, which can only bind to tau monomers.

### Chaperone interactions with tau

Overall it appears that the DNAJB1 and DNAJA2 chaperones, despite being very similar in structure, display distinct differences in their interaction with tau. While DNAJA2 binds to all tau species - monomers, aggregation-prone tau, and fibers - DNAJB1 does not bind at all to inert tau monomers, but interacts strongly with the aggregation-prone species as well as mature fibers.

This disparity between the chaperones could be explained by utilization of different structural domains to recognize the various tau species, however no structural information is currently available for either DNAJA2 or DNAJB1 in complex with tau^4R^. We therefore utilized NMR to map the binding sites on the chaperones for the various tau species.

Both DNAJA2 and DNAJB1 are homodimeric proteins comprised of an eponymous N-terminal J-domain (JD) that is essential for Hsp70 activation, two putative substrate binding domains, CTDI and CTDII, and a C-terminal dimerization domain. In addition, DNAJA2, as all class A J-domain proteins, has a zinc-finger like region (ZFLR) insertion in CTDI (40–42), while DNAJB1 has an autoinhibitory GF region connecting the J-domain to CTDI and blocking premature interaction of Hsp70 with the JD (43).

Due to the large size of the DNAJA2 dimer (90 kDa), which hampers NMR experiments, we used a monomeric version of the protein that contained only the substrate-binding and ZFLR domains (44). This construct (27 kDa) of ^2^H, ^15^N-labeled DNAJA2^111-351^ was far more amenable to NMR and gave a high quality ^1^H-^15^N HSQC-TROSY spectrum (41) (Figure S7A). Addition of two-fold excess of tau^4R^ caused CSPs in the first DNAJA2 substrate-binding domain (44), located in CTDI (Fig. 6A, C and S7A). Specifically, tau^4R^ (most likely via the hydrophobic PHF6 and PHF6* motifs) binds to a hydrophobic pocket located between β-strands 1 and 2 (Figure 6C, colored purple). The binding was in fast exchange and no reduction in peak intensities to DNAJA2 residues was observed (Fig. S7A), in agreement with the relatively low affinity of DNAJA2 for monomeric tau (43 μM, Fig 2C). DNAJB1, despite its CTDI having 56% identity to the CTDI tau-binding region of DNAJA2, showed no interaction with the monomeric tau (Fig. 6E and S7C and D), as previously seen from the tau side (Fig. 1C).

**Figure 6.**
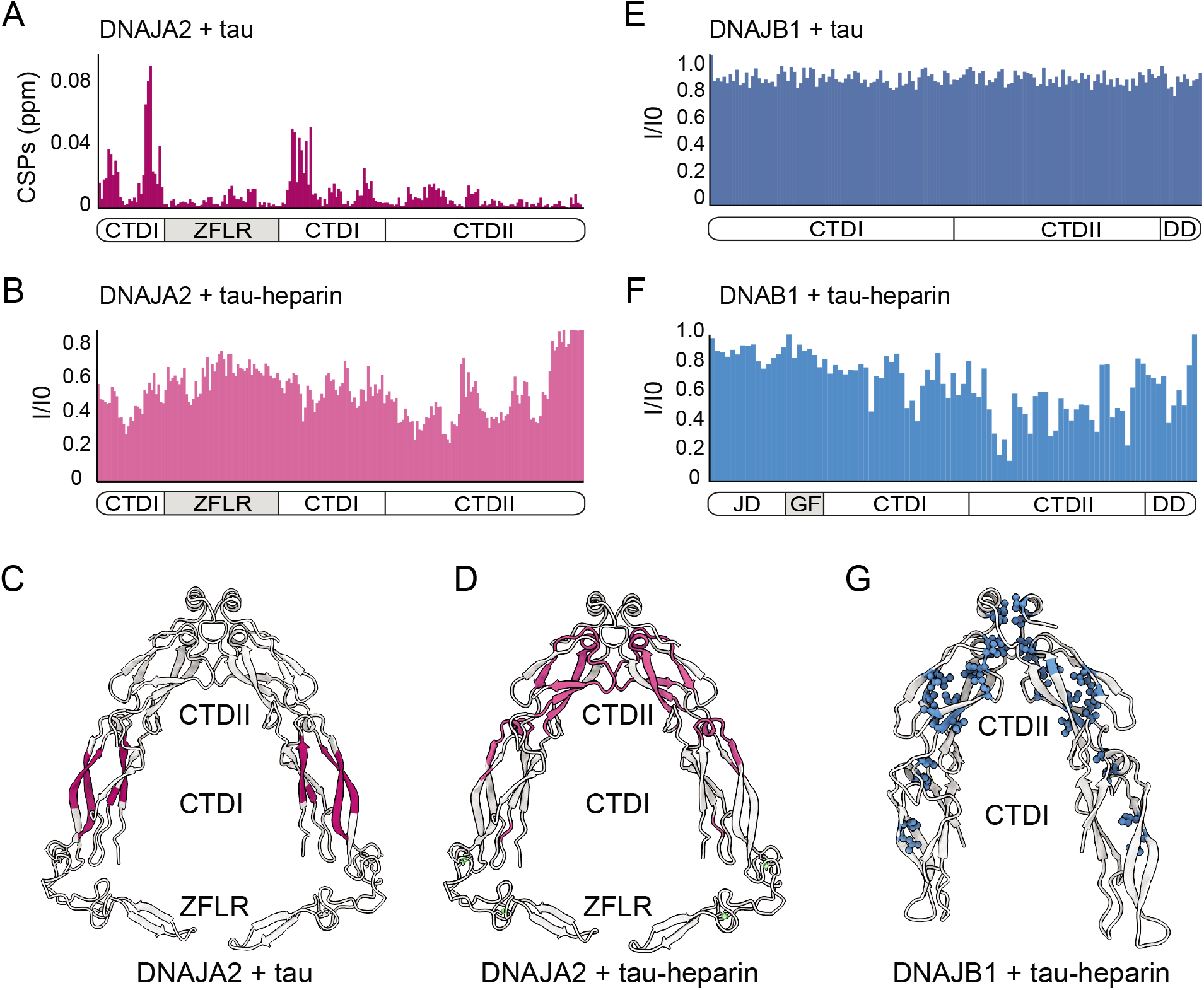
Mapping DNAJA2 and DNAJB1 chaperone binding to tau species. **(A)** Combined amide chemical shift perturbations between free DNAJA2 and DNAJA2 bound to unlabeled tau. The domain organization of DNAJA2 is indicated at the bottom. Monomeric tau binds to the C-terminal domain I (CTDI) of DNAJA2, but not to the ZFLR insertion. **(B)** Residue-resolved backbone amide NMR signal attenuation (I/I_0_) plot for DNAJA2 in complex with aggregation-prone tau species, generated by addition of two-fold excess heparin. The binding is primarily to the C-terminal domain II (CTDII) of DNAJA2. **(C-D)** Structural representation of DNAJA2 chaperone, with residues showing significant CSPs upon binding to monomeric tau (from panel A) highlighted in purple (C), and residues displaying significant decreases in peak intensities upon binding to aggregation-prone tau species colored pink (D). **(E)** No changes in signal intensity are detected in the CTDs of DNAJB1 upon addition of 2-fold excess of monomeric tau protein, indicating a lack of interaction. **(F)** Combined methyl group chemical shift perturbations between free DNAJB1 and DNAJB1 bound to aggregation-prone tau species generated by addition of two-fold excess heparin. Binding is observed to the CTDII domain of DNAJB1. **(G)** Structural representation of DNAJB1 chaperone with methyl residues showing significant changes upon binding to aggregation-prone tau species highlighted in blue.

We then repeated the binding experiment using aggregation-prone tau species, generated by the addition of heparin (35). Interestingly, this tau species, unlike the inert tau monomer, caused significant peak broadening to residues located in the second substrate binding region of CTDII of DNAJA2 (Fig. 6B and S7B), whereas only small changes were detected in the CTDI region (Fig. S7B). Hence, the inert monomeric tau interacts predominantly with the DNAJA2 substrate-binding groove in CTDI, while the aggregation-prone tau species preferentially binds to CTDII (Fig. 5C and D). Having a second, distinct tau-binding domain thus explains both the high affinity of DNAJA2 towards the aggregation-prone tau, and the ability of the chaperone to interact with the two tau species.

We then tested the binding of DNAJB1^151-341^ to this aggregation-prone tau species. The interaction with tau-heparin, however, caused severe peak broadening, preventing us from obtaining site-specific information. The high molecular mass of the complex formed between multiple aggregation-prone tau monomers and the DNAJB1-dimer is presumably responsible for this effect. To overcome this problem we utilized a ^2^H ^13^CH_3_-ILVM sample of full-length DNAJB1 that gave a high quality ^1^H-^13^C HMQC spectrum even upon complex formation with the aggregation-prone tau (Fig. S7E). Selective peak broadening was detected in methyl residues located in CTDII (Fig. 5F, G and S7E) – indicating that, similarly to DNAJA2 chaperone, the aggregation-prone tau binds to CTDII in DNAJB1. This CTDII site in DNAJB1 was also recently identified as a binding site for another amyloid-forming protein, α-synuclein (43).

Hence, DNAJB1 and DNAJA2 recognize seed-competent tau^4R^ species predominantly via CTDII, yet, the structural differences between the CTDI domains of the chaperones likely cause the diverging specificities for the various tau^4R^ species.

## DISCUSSION

Hsp70 chaperones are known to be key factors in tau quality control and turnover (45), however the contributions of their Hsp40 co-chaperones remain poorly understood. In this study, we describe the effects of Hsp40 chaperone family members, DNAJA2 and DNAJB1, on tau amyloid fiber formation in comparison to the well-characterized tau aggregation suppressor HSPB1.

DNAJA2 chaperone was previously identified as a potent suppressor of tau aggregation (13). Our results demonstrate that DNAJA2 can interact simultaneously with multiple tau species, which can explain its high effectiveness in aggregation prevention. DNAJA2 does not only bind tau monomers via its CTDI, thus preventing fiber growth by monomer addition, but it also binds aggregation-prone tau species via its CTDII, which effectively reduces the speed of nucleation. Moreover, DNAJA2 even associates with mature tau fibers, which provides an additional pathway for inhibiting the incorporation of new monomers into the growing fibers.

Given this multiplicity of interfering pathways, DNAJA2 has the potential to serve as an early protective cellular factor that limits tau aggregation, which explains the correlation between distorted cellular DNAJA2 levels and pathology-linked nucleation sites (13).

Remarkably, in comparison to DNAJA2, DNAJB1 displayed significantly different interactions with tau despite the high structural similarity between both chaperones. In fact, unlike DNAJA2, DNAJB1 does not interact with inert tau monomers, although its CTDI is highly homologous to that of the class A chaperone. DNAJB1 affinity however, is increased by orders of magnitude for the aggregation-prone tau species. There, DNAJB1 also functions as a bona fide chaperone, effectively hindering tau aggregation by preventing the formation of the seeding nuclei, as well as by stably binding to the amyloid fibers themselves, slowing down their growth. Interestingly, both interactions are mediated by CTDII, thus leaving the CTDI site free to potentially recruit Hsp70 chaperones and initiate the disaggregation of fibers when these are formed.

This mode of action of DNAJB1 differs significantly from that of the well characterized tau aggregation-suppressor HSPB1, as well as from the recently described activity of the DNAJC7 chaperone (46). These chaperones preferentially bind to the PHF6 repeats in inert tau monomers, thus protecting these aggregation-prone regions and preventing them from being incorporated into the growing fibers. Moreover, by solely interacting with inert tau monomers, HSPB1 can only effectively inhibit the rate of fibril elongation, but cannot directly affect the final length of the tau amyloids.

In contrast, the actions of DNAJA2 and DNAJB1 result in a significant decrease not only in fibril mass, but also length. This function of the Hsp40 chaperones, compared to HSPB1, could be attributed to their ability to interact with the forming tau fibers, which efficiently blocks the incorporation of additional tau monomers. In addition, the Hsp40 chaperones bind aggregation-prone tau species, thereby blocking their elongation into the mature tau fibers.

The relatively minor differences in the aggregation prevention abilities of DNAJB1 and DNAJA2, along with the relatively poor performance of HSPB1 in preventing fibril growth, suggest that, while interaction with soluble tau monomers helps slow aggregation, this does not significantly contribute to the ability of chaperones to prevent amyloid growth once it has begun. It is therefore possible to envision a scenario in which HSPB1 delays amyloid formation during the early stages of tau aggregation via interaction with the monomers, and Hsp40 chaperones later interact with seeds and more mature species to further hinder the fibril formation process.

One open question is how these chaperones affect the aggregation of tau mutants linked to tauopathies. It has been hypothesized that some variants, such as for example P301L, may be capable of “avoiding” the chaperone system, thus possibly contributing to the disease pathology. Such behavior was indeed recently observed with the DNAJC7 chaperone, which was found to have a significantly reduced affinity to the tau P301L mutation (46). Furthermore, in our aggregation prevention assays, a reduction was observed in the activity of HSPB1 when incubated with the P301L variant compared to wild-type tau (13) (Fig. S6B). For these two chaperones, their reduced efficacy is likely due to the equilibrium of the P301L mutation shifting towards an aggregation-prone seeding conformation of tau (34), for which both DNAJC7 (46) and HSPB1 (Fig. 4B) display greatly reduced affinities. In contrast, DNAJA2 and DNAJB1 remained effective in suppressing the aggregation of the P301L variant of tau^4R^ (with the anti-aggregation activity of DNAJB1 being even higher for this tauopathy mutant, Fig. S6C), indicating that these Hsp40 chaperones could be effective in suppressing a wide range of tauopathies.

A second, crucial question that has yet to be answered is whether the reduction of fibral growth by the Hsp40 chaperones, which results in generation of smaller tau fibrils, is indeed beneficial for cellular homeostasis. A recent study showed that the disaggregation of tau by the DNAJB1/Hsp70/Hsp110 chaperones generates low-molecular-weight tau species, which were seeding-competent in cell culture models (23). Hence, chaperone-mediated tau disaggregation may not be beneficial per se, but may instead be involved in the prion-like propagation of tau pathology. The smaller tau species generated during the DNAJA2 and DNAJB1 aggregation-prevention processes could then have similar prion-like propagation properties, acting as seeds that can sequester more tau protein into amyloid aggregates. In such a case, chaperone-mediated aggregation-prevention would, in fact, accelerate the progression of disease, ultimately proving detrimental to cell health. HSPB1 chaperones, on the other hand, do not generate smaller tau species during their aggregation prevention, and could therefore be more beneficial in slowing the progression of disease, despite their lower chaperoning activity.

However, another important aspect to consider is that aggregation prevention in the cell does not occur in isolation and can also be coupled to protein degradation via the proteasome or autophagy. These pathways could potentially be more potent in degrading smaller tau fibrils and aggregation-prone monomers than the mature fibrils, thereby rendering the activity of DNAJA2 and DNAJB1 beneficial overall.

Hence, further studies will be required to understand the full role of Hsp40-mediated tau aggregation prevention in the cell, as well as to evaluate the therapeutic potential of the Hsp40 chaperone machineries in combating tauopathies.

## MATERIALS AND METHODS

### Construct Preparation

Tau^4R^ (residues 244-372, C322A) wt and mutants, HSPB1 (S15D, S78D, S82D, I181G, V183G), DNAJA2, and DNAJB1 (residues 154-341) were expressed in *E. coli* BL21 (DE3) cells from pET-29b(+) vector with a N-terminal His_6_ tag followed by a tobacco etch virus (TEV) protease cleavage site. Tau (C322S), DNAJB1, and Ydj1 (residues 111-351, F335D; yeast orthologue of DnaJA), were expressed from the pET-SUMO vector with an N-terminal His_6_ purification tag and an Ulp1 cleavage site (DNAJB1 plasmid was a gift from B. Bukau, University of Heidelberg).

### Protein expression

Cells were grown in Luria Bertani broth (LB) to OD_600_≈0.8 at 37 °C and expression was induced by addition of 1 mM isopropyl-β-D-thiogalactoside (IPTG). Cells expressing HSPB1, DNAJA2, and DNAJB1 chaperone variants were allowed to proceed overnight at 25 °C and cells expressing tau constructs at 18 °C.

Isotopically labeled tau and tau^4R^ proteins for NMR were grown in M9 H_2_O media supplemented with ^15^NH_4_Cl (and ^13^C-glucose) as the sole nitrogen (and carbon) source. Protein expression was induced with 1mM IPTG at 18 °C overnight.

Labeled DNAJB1^154-341^ and Ydj1^111-351^ were grown at 37 °C in M9 D_2_O media supplemented with [^2^H,^12^C]-glucose and ^15^NH_4_Cl as the sole source of carbon and nitrogen. In the case of DNAJB1, 2-ketobutyric acid-^13^C_4_,3,3-d_2_ sodium salt (60 mg/L), 2-ketoisovaleric acid-^13^C_4_,d_3_ sodium salt (80 mg/L), and ^13^C-L-methionine (100 mg/L) (Cambridge Isotope Laboratories) were added 1 hour prior to induction with 1mM IPTG, following the procedure of Tugarinov *et al*. (47) to produce U-^2^H, ^15^N, ^13^CH_3_-ILVM labelled protein. Proteins were expressed at 25 °C overnight.

### Purification of labeled and unlabeled proteins

Proteins were purified on a Ni-NTA HiTrap HP column (GE Life Sciences). The purification tag was cleaved by the appropriate protease (see Construct Preparation) and the cleaved protein was further separated from the uncleaved protein, the tag, and the protease on a Ni-NTA HiTrap HP column. HSPB1, DNAJA2, and DNAJB1 chaperone variants were concentrated on an Amicon Ultra-15 10K molecular weight cutoff (MWCO) filter (Millipore) and further purified on a HiLoad 16/600 Superdex 200 pg gel filtration column (GE Healthcare), equilibrated with 25 mM HEPES pH 7.0, 150 mM KCl, and 2 mM DTT. Tau constructs were concentrated on an Amicon Ultra-15 3.5K MWCO filter (Millipore) and further purified on a HiLoad 16/600 Superdex 75 pg gel filtration column (GE Healthcare) equilibrated with 25 mM HEPES pH 7.0, 300 mM KCl, and 2 mM DTT. Purity of proteins was confirmed by SDS-PAGE.

### Aggregation prevention assays

Aggregation kinetics were measured in Synergy H1 microplate reader (BioTek) in black, flat-bottom, 96-well plates (Nunc).Tau or tau^4R^ variants (10 μM) were pre-incubated in the presence or absence of indicated chaperones for 10 minutes at 37 °C. All proteins in the assay were buffer exchanged into the assay buffer (50 mM HEPES pH 7.4, 50 mM KCl, and 2 mM DTT). Thioflavin T (ThT; Sigma) at a final concentration of 10 μM was added and the aggregation was induced by the addition of a freshly prepared heparin salt solution (Sigma). Aggregation reactions were run at 37 °C with continues shaking (567 rpm) and monitored by ThT fluorescence (excitation = 440 nm, emission= 485 nm, bandwidth), using an area scan mode with a 3×3 matrix for each well. Black, flat-bottom, 96-well plates (Nunc) sealed with optical adhesive film (Applied Biosystems) were used. For data processing, baseline curves at same conditions but without heparin were subtracted from the data. Samples were run in triplicate and the experiments were repeated at least 4 times with similar results.

### Seeded tau aggregation reactions

Tau seeds were prepared from mature tau fibers generated under similar conditions to these in the aggregation prevention assays, except that ThT was omitted. The fibers were then sonicated using a probe sonicator (Vibra-Cell, SONICS) with an amplitude of 40%, for 30 seconds on and 10 seconds off, for a total of 7 minutes. The sonicated fibers were immediately added to monomeric tau, ThT and DTT in a 96-well plate in the ratios described above and ThT fluorescence was measured as a function of time.

### Dynamic light scattering (DLS)

The hydrodynamic radius of tau^4R^ seeds was measured by dynamic light scattering on a DynaPro DLS Plate Reader III (Wyatt Technology). Tau seeds (10 μM) were loaded on a 96-well black, clear bottom plates (Nunc) and subjected to a 5-min 3,000 × g centrifugation to remove air bubbles from the wells. Measurements were carried out 20 times per well before averaging, with 5 second acquisitions at 25 °C. Resulting autocorrelation functions were fitted with the equation

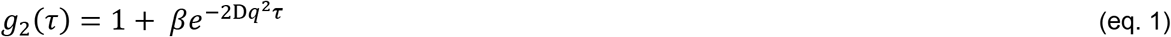

where *β* is the coherence factor, D is the translational diffusion coefficient, and q is the scattering wave vector given by

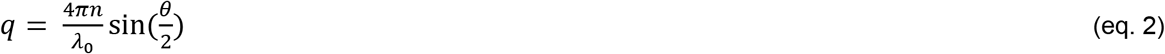

where n is the solvent refractive index (n = 1.334 was used), *λ*_0_ is the wavelength used by the instrument, and *θ* is the scattering angle.

The Stokes-radius (*R_s_*) was calculated from the translational diffusion coefficient, *D*, using the Stokes–Einstein equation

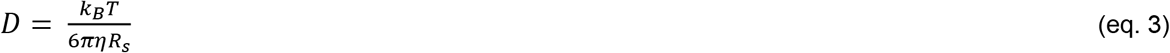

Where k_B_ is Boltzmann coefficient 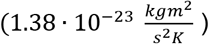, T the temperature (298 K), and *η* is the viscosity of our buffer (1.03).

### Aggregation Prevention Data Fitting

All aggregation kinetics were fitted with a saturation-elongation-fragmentation model (31) using a critical nucleus size of *n_c_* = 2. The differential equation system for this model is -

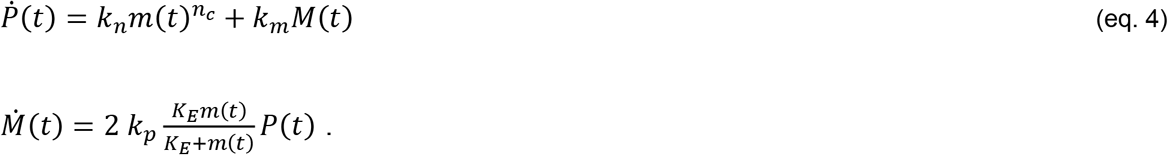

Here, *P*(*t*) is the number concentration of fibrils, *M*(*t*) is the mass-concentration of a fibril, *m*(*t*) is the monomer concentration, *k_n_* is the nucleation rate, *k_m_* is the fragmentation rate, *k_p_* is the elongation rate, and *K_E_* is the equilibrium constant for monomer addition to an existing fibril. For fitting, the ThT-fluorescence signal *S_i_*(*t*) of the i^th^ time trace was converted to the mass-concentration of the fibrils *M_i_*(*t*) according to

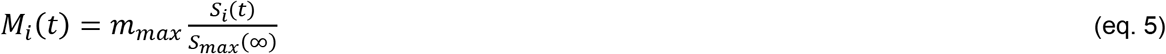

Here, *m_max_* is the highest tau^4R^ concentration used in the experiments and *S_max_*(∞) is the ThT-signal of the long-term plateau for the time trace with the highest tau^4R^ concentration. The resulting mass concentration *M_i_*(*t*) was then fitted by numerically solving the differential equation system eq. 4 for *M*(*t*), using the initial conditions *M*(0) = 0, *P*(0) = 0, and *m*(0) = *m_i_*, where *m_i_* is the initial concentration of tau^4R^ monomers for the i^th^ time trace. Prior to fitting, the time traces *M_i_*(*t*) were smoothed by binning data points to reduce noise and to speed up fitting. The bin size was 2 – 5 data points. Fitting was performed using the “Differential Evolution” method (48) in Mathematica 11.2 (Wolfram). Importantly, *k_p_* cannot be independently obtained from unseeded data. We therefore arbitrarily set *k_p_* = 1 for the global fit of the unseeded data at all monomer concentrations, thus obtaining *k′_n_* = *k_n_k_p_* and *k′_m_* = *k_m_k_p_*. In a second step, we determined *k_n_*, *k_m_*, and *k_p_* in a global fit of a data set including aggregation seeds using

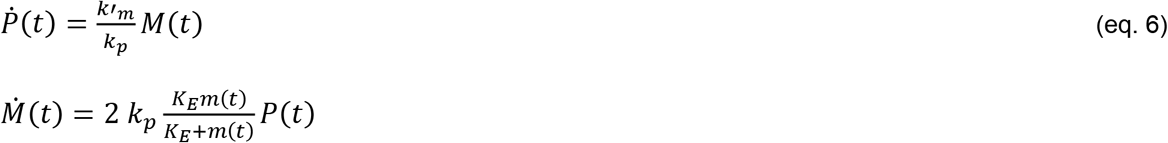

with the initial conditions *M*(0) = *M*_0_, *P*(0) = *M*_0_/*L*, and *m*(0) = *m*_0_ = 10 *μM*. Here, *M*_0_ is the mass concentration of the seeds and *L* is the length of the seeds. We determined *L* by measuring the Stokes-radius *R_s_* = 55 *nm* of the seeds using dynamic light scattering (DLS, see above). We then modeled the seeds as an ellipsoid with a long axis *a* (seed length) and a short axis *b* (fibril thickness). The friction coefficient for an ellipsoid is given by *ξ_e_* = 6*πηa*/ln(2*a/b*), which must be identical to that of a sphere with the Stokes-radius *R_s_* given by *ξ_s_* = 6*πηR_s_*. From the equality, the fibril length *a* can be determined given that the fibril thickness *b* is known. Based on existing cryo-EM structures of tau-fibrils (6QJH, 6QJM, 6QJP), we estimated *b*~10 *nm*, which results in *a*~200 *nm*. Given the spacing of tau monomers in a fibril of approximately 2 nm, we estimated a seed length of *L* = 100 tau monomers. Error estimates of the kinetic rates were obtained by fitting two independent data sets.

Fitting of the data in the presence of chaperones were performed for each kinetic trace individually by fixing the kinetic rates to those determined in the absence of chaperone and only allowing one rate to vary at a time. An exception was the data set for DNAJA2 in which we allowed the simultaneous variation of *k_n_* and *k_p_*. Since only a moderate parameter space was scanned, we used “Simulated Annealing” to optimize the parameters. We would like to note, that the amplitude of the aggregation kinetics was not a free fitting parameter but was determined by the total concentration of tau^4R^ monomers (10 μM) in the experiment. Hence, fitting of the aggregation kinetics in the presence of chaperones assumes that the presence of chaperone does not alter the ThT-concentration accessible in solution to stain the fibrils.

### NMR Spectroscopy

All NMR experiments were carried out at 25 °C on 14.1T (600 MHz), 18.8T (800 MHz), or 23.5T (1000 MHz) Bruker spectrometers equipped with triple resonance single (z) or triple (x,y,z) gradient cryoprobes. The experiments were processed with NMRPipe (49) and analyzed with NMRFAM-SPARKY (50) and CCPN (51).

### NMR Assignment experiments

Assignments for tau^4R^ were transferred from the BMRB (entry 19253) and corroborated by HNCACB, CBCA(CO)NH, HN(CA)CO, and HNCO experiments on a 4 mM sample of [U-^15^N,^13^C]-labeled tau^4R^ in 50 mM HEPES pH 7.4, 50 mM KCl, 1 mM DTT, 0.03% NaN_3_, and 10% D_2_O. The assignment experiments were recorded on an 800 MHz magnet, resulting in the unambiguous assignment of 88% of non-proline residues.

Tau-heparin complex assignments were obtained by recording 3D HNCA, CBCA(CO)NH, and HN(CA)CO on a 2.5 mM [U-^13^C,^15^N]-labeled tau^4R^ sample supplemented with 2.5 mM heparin. The experiments were recorded on an 800 MHz magnet and 84% of non-proline residues were assigned.

### Secondary Structure Propensities

Secondary structure propensities for tau4R and tau4R-heparin complex were calculated from backbone C’, C_α_, and C_β_, 1H, 15N chemical shifts following a procedure described in Marsh et al. (52). Proline and cysteine residues were omitted from this calculation.

### NMR tau-chaperone binding experiments

Tau interaction with chaperones was assayed for 200 μM samples of [U-^15^N]-labeled tau, tau^4R^, or tau^4R^ P301S and P301L mutants. Tau variants were measured alone or upon addition of heparin (200 μM) and/ or chaperones (100 or 400 μM; as indicated in spectrum) in 50 mM HEPES pH 7.0, 50 mM KCl, 1 mM DTT, 0.03% NaN_3_, and 10% D_2_O. ^1^H-^15^N HSQC-TROSY spectra were acquired for each sample and peak intensities were determined by quantifying peak volumes. Regions of tau^4R^ with signal loss greater than one standard deviation from the average intensity ratio were determined to be the regions of binding.

High salt binding experiments were performed with samples of ^15^N-tau^4R^ (200 μM) and 400 μM [U-^1^H]-labeled chaperones in 50 mM HEPES pH 7.0, 300 mM KCl, 1 mM DTT, 0.03% NaN_3_, and 10% D_2_O. Binding was determined by calculating intensity ratios as described above.

The binding of DNAJB1 and DNAJA2 chaperones to tau^4R^ was measured by acquiring ^1^H-^15^N HSQC-TROSY spectra for 200uM [U-^2^H,^15^N]-labeled DNAJB1^154-341^ or Ydj1^111-351^ alone or with 2-fold excess of deuterated tau^4R^. The reactions were measured in 50 mM HEPES pH 7.0, 50 mM KCl, 1 mM DTT, 0.03% NaN_3_, and 10% D_2_O. Backbone DNAJB1^154-341^ and Ydj1^111-351^ assignments were available through the BMRB (entries 27998 and 28000, respectively).

The interaction of full length DNAJB1 to tau^4R^ was determined by acquiring ^1^H-^13^C HMQC methyl-TROSY spectra (53) for 100uM [^2^H, ^13^CH_3_]-ILVM labeled DNAJB1 alone or with 200 μM ^2^H-tau^4R^ (or tau-heparin complex) in 50 mM HEPES pH 7.4, 100 mM KCl, 2 mM DTT and 0.03% NaN_3_ in 100% D_2_O. ILVM assignments for full length DNAJB1 were taken from previous work in our lab (43). Binding regions were determined by intensity ratio as described above.

### NMR chemical shift perturbations

The interaction of tau^4R^ with heparin was monitored by 2D ^1^H–^15^N HSQC experiments. Heparin (40-400 μM) was titrated into 200 μM of ^15^N-labeled tau^4R^) in 50 mM HEPES pH 7.0, 50 mM KCl, 1 mM DTT, 0.03% NaN_3_, and 10% D_2_O and chemical shifts were recorded.

CSPs were calculated from the relation

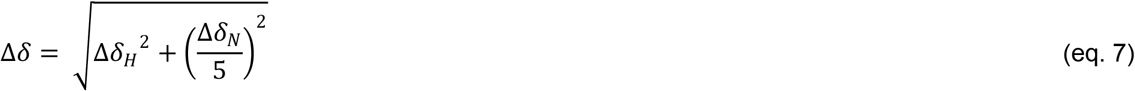

where Δδ_H_ is the amide proton chemical shift difference and Δδ_N_ is the ^15^N backbone chemical shift difference. CSPs greater than one standard deviation from the mean were considered significant.

### Negative stain electron microscopy

Tau fibrils or tau fibrils-chaperone mixtures (10 μl) were deposited on glow discharged carbon-coated copper EM grids (Electron Microscopy Sciences), washed with three consecutive drops of 1% w/v Uranylformate, and air-dried. Imaging was performed on an FEI T12 Spirit transmission electron microscope at 120kV and a magnification of 9300-30000 times, equipped with a Gatan OneView CMOS 4K x 4K CCD camera.

### Chaperone-fibril co-sedimentation assay

Preformed tau^4R^ fibers (10 μM) were incubated with HSPB1, DNAJB1, and DNAJA2 chaperones (10 μM) for 20 min at 37 °C in 50 mM HEPES pH 7.4 and 50 mM KCl. Tau fibers were separated from the unbound chaperones by centrifugation at 16,900 g for 30 minutes. The pellets were washed, resuspended in 50 μL of buffer with 20% SDS, and sonicated for 10 minutes. Samples were incubated for 5 minutes at 95 °C and run on a 4-20% gradient SDS-PAGE gel (Genscript).

### Fluorescence anisotropy measurements

Steady-state equilibrium binding of DNAJA2, DNAJB1 chaperones to preformed tau fibers was measured by fluorescence polarization using 100 nM of fluorescently tagged chaperones (DNAJB1 G194C-AF488 or DNAJA2-AF488).

Steady-state equilibrium binding of DNAJB1, DNAJA2 and HSPB1 to monomeric tau was measured by fluorescence anisotropy using 100 nM of fluorescently tagged tau (tau C291S, C322S, L243C-AF488). Samples were allowed to equilibrate for 10 minutes at 37 °C and measurements were performed on a Tecan SPARK 10M plate reader in black, flat-bottomed 384 square well plates. Measurements were performed on a Tecan SPARK 10M plate reader in black, flat-bottomed 384 square well plates. The excitation filter was centered on 485 nm with a bandwidth of 20 nm, and the emission filter was centered on 535 nm with a bandwidth of 25 nm. The gain and Z position were optimized from a well in the center of the binding curve, followed by calibration of the G factor. 60 flashes were performed per well.

Data were fit to a one-site binding model using OriginPro version 2018.

